# Kappa opioid receptors (KORs) in the anterior paraventricular nucleus of the thalamus (aPVT) mediate morphine withdrawal-, anxiety-, fear-, and KOR agonist-induced aversion-like behaviors in mice

**DOI:** 10.64898/2026.06.30.735625

**Authors:** Peng Huang, Chongguang Chen, Kathryn Bland, Aryan Anand, Kevin Beier, Lee-Yuan Liu-Chen

**Affiliations:** Center for Substance Abuse Research and Department of Neural Sciences, Temple University Lewis Katz School of Medicine, Philadelphia, PA; Department of Physiology and Biophysics, School of Medicine, University of California, Irvine, CA

## Abstract

PVT is involved in stress responses, fear, anxiety, arousal, aversion and reward. Anterior and posterior PVT (aPVT and pPVT) have different neuronal connections, molecular contents, and functional roles. PVT expressed a high level of KOR. Herein we mapped the projection targets of aPVT KOR(+) neurons and explored the behavioral significance of aPVT KOR. Using KOR-iCre mice and Cre-dependent anterograde tracer, we found that KOR(+) glutamatergic neurons in aPVT primarily projected to NAc, CeA, BNST, PFC, and reticular nucleus of the thalamus (RT). 3-D images showed the pathway emanating from aPVT to the ventral RT, through NAc, and out to the other regions, indicating widespread axonal collateralization. We conditionally knock-downed KOR (KOR cKD) in aPVT by injection of AAV-eGFP-Cre or AAV-eGFP (control) into aPVT of Oprk1^lox/lox^ mice. [^3^H]U69,593 receptor autoradiography revealed substantial KOR cKD in aPVT. In both male and female mice, the KOR cKD in aPVT significantly reduced anxiety-like behaviors in the elevated plus-maze test, cue-induced freezing after fear conditioning and naloxone-precipitated morphine withdrawal-associated jumps. KOR cKD attenuated U50,488H-induced conditioned place aversion in males only, while having no effect on forced swim immobility or the U50,488H-induced visceral analgesic and antipruritic effects in either sex. Thus, our results reveal for the first time that KOR-mediated inhibition of aPVT neurons may mediate morphine withdrawal, anxiety, and cue-induced fear in both sexes but contribute to KOR agonist-induced aversion only in males. Notably, our findings reveal a previously unrecognized role for aPVT in regulating morphine withdrawal, acting in a manner distinct from pPVT.

## INTRODUCTION

The kappa opioid receptor (KOR), one of the three opioid receptors, is a G_i/o_ protein-coupled receptor. Activation of KOR produces many effects, including analgesic and anti-pruritic effects and water diuresis [1–3]; however, it also produces side effects, including dysphoria, psychotomimesis, and sedation [4–7]. KOR also plays an important role in stress-induced responses and pro-addictive behaviors and is implicated in anxiety and depression (see [8–12] for reviews). The neuronal bases for these actions are not fully elucidated.

The paraventricular nucleus of the thalamus (PVT) extends over the entire rostro-caudal length of the most dorsal midline thalamus. It receives convergent inputs from several cortical regions, including the prelimbic, infralimbic, and insular cortices, as well as from the ventral subiculum, hypothalamic nuclei, and brainstem. It, in turn, projects densely to multiple limbic and forebrain targets, notably the nucleus accumbens (NAc), central amygdala (CeA), bed nucleus of the stria terminalis (BNST), hypothalamic nuclei, and prefrontal cortex (PFC) [reviewed in [13–15]]. Notably, tracing studies of neurons that primarily project to individual targets such as the nucleus accumbens shell (NAcSh), dorsolateral BNST (BSTDL), central amygdala lateral division (CeL), infralimbic cortex, or the dorsomedial hypothalamic nucleus revealed widespread axonal collateralization [16, 17]. These neurons exhibited fiber labeling across all major PVT-innervated regions, albeit with varying densities, indicating that PVT projection neurons can influence multiple limbic circuits simultaneously [17]. Consistent with these anatomical features, the PVT has been increasingly implicated in the regulation of behavioral responses related to stress, fear, anxiety, arousal, feeding, motivation, and reward [reviewed in [13, 14, 18]]. It is common to divide the PVT into anterior PVT (aPVT) and posterior PVT (pPVT) due to differences in afferent and efferent projections, neurotransmitters, and roles in behaviors. The aPVT has been linked to appetitive/approach behaviors, whereas the pPVT is associated with aversive/avoidance behaviors (reviewed in [18, 19]). Behavioral studies have historically emphasized the pPVT [14], while research on the aPVT has recently begun to expand (e.g., [20–32]). In mice, the PVT spans ∼ 2.1 mm along the rostro-caudal axis. The division between aPVT and pPVT in the literature is somewhat arbitrary. In this study, we used the following coordinates for aPVT viral injections (relative to the bregma): AP - 0.50, ML 0.0, DV −3.40 according to Paxinos and Franklin [33].

In mice and rats, the PVT exhibits robust expression of KOR mRNA, as revealed by *in situ* hybridization (ISH) ([34–36] also Allen Brain Institute’s Mouse Brain ISH Atlas) and single nucleus RNA sequencing (snRNA-seq) [37]. High level of KOR protein was also detected in PVT by autoradiography of radioligand binding in mice and rats [34, 35] and KOR conjugated in frame with tdTomato [38]. Based on a recent snRNA-sequence analysis of PVT neurons in mice, KOR mRNA expression was found to be more abundant in aPVT compared with pPVT [37]. In accord with this KOR distribution difference, dynorphin A elicited prominent inhibitory outward K⁺ currents in aPVT neurons, whereas such responses were weaker or less frequent in pPVT neurons in mice [39]. Across mouse studies, activation of KOR in the PVT consistently inhibited PVT neuronal cell bodies by inducing postsynaptic hyperpolarization mediated by outward K⁺ currents [39, 40]. In the PVT, KOR activation has also been shown to reduce synaptic input through presynaptic mechanisms. In rat pPVT slices, dynorphin A decreased the frequency of spontaneous and miniature excitatory postsynaptic currents without affecting amplitude, indicating a presynaptic reduction in glutamate release onto PVT neurons [41]. Complementing this, mouse PVT neurons co-expressing KOR and μ-opioid receptors (MOR) exhibited reduced glutamatergic and GABAergic synaptic inputs upon KOR activation, consistent with presynaptic modulation of both excitatory and inhibitory afferents [40]. In addition, in mice, KOR regulated the inhibitory input from the zona incerta (ZI) to PVT, and chronic morphine exposure removed this inhibition (disinhibited) this pathway [42]. These presynaptic effects occur alongside the postsynaptic hyperpolarization [39, 40], revealing a dual mechanism by which KOR regulates PVT activity. Collectively, these studies demonstrate that KORs in the PVT act both post-synaptically and pre-synaptically within PVT to shape the thalamic output.

In this study, we mapped the projections of the postsynaptic KOR(+) neurons in aPVT by anterograde tracing and examined the role of KOR in aPVT neurons in several mouse behaviors associated with PVT function and KOR signaling by selectively deleting KOR from aPVT neurons.

## MATERIALS AND METHODS

### Drugs

U50 (U50,488H; U50,488 methanesulfonate) and morphine sulfate were obtained from the National Institute on Drug Abuse Drug Supply Program (Bethesda, MD). Tamoxifen, naloxone hydrochloride, Compound 48/80, acetic acid, and paraformaldehyde were purchased from Sigma-Aldrich. Unless otherwise indicated, drugs or vehicle (saline) were administered subcutaneously (s.c.) at 10 µl/g body weight. Compound 48/80 (0.5 mg/ml) was injected into the nape at 100 µl (50 µg) / mouse, while acetic acid (0.6%, v/v) was administered intraperitoneally (i.p.) at 10 µl/g body weight. Mice received tamoxifen (3 mg / mouse) by oral gavage once daily for 5 consecutive days to induce Cre expression. [^3^H]U69,593 (46 Ci/mmol) was purchased from Revvity (Waltham, MA).

### Viral vectors

For anterograde tracing, Cre-dependent AAV1-FLExloxP-mGFP-2A-synaptophysin-mRuby was provided by Dr. Kevin T Beier of University of California, Irvine. In addition, AAV8-hSyn-DIO-mCherry was purchased from Addgene (50459) (Watertown, MA). For conditioned KOR knockdown, AAV9-eGFP-Cre and AAV9-eGFP were purchased from Addgene (105540-AAV9/pENN.AAV.hSyn.HI.eGFP-Cre.WPRE.SV40 and 50465-AAV9/pAAV-hSyn-EGFP, respectively).

### Mice

#### Adult male and female mice were used for all the experiments

Homozygous Oprk1^lox/lox^ mice (B6;129-*Oprk1^tm2.1Kff^*/J, JAX:030076) were obtained from the Jackson Laboratory (Bar Harbor, ME). This strain of mice, originally generated by the laboratory of Brigitte Kieffer (Strasbourg University, France) and possess *loxP* sites flanking exon 3 of the KOR gene (*Oprk1*). KtdT is a mutant mouse line expressing the KOR conjugated in frame at the C-terminus with the fluorescent protein tdTomato (KOR-tdT or KtdT). KtdT mice were custom generated by Cyagen Co. (Santa Clara, CA) for our laboratory as we described previously [38] and cryopreserved by the Jackson Laboratory as *Oprk1tm1.1Llch*. KOR-iCre is another mutant mouse line custom generated by Cyagen Co. for our laboratory that expresses KOR linked to CreERT2 via a 2A peptide. The KOR-iCre mice were cryopreserved by the Jackson Laboratory as *Oprk1tm2.1(icre/ERT2)Llch*. The targeting strategy for this line is illustrated in Supplemental Figure 1. Both KtdT and KOR-iCre mouse lines were generated and maintained on a pure C57BL/6N background.

#### Wildtype C57BL/6N mice were purchased from the Jackson Laboratory

In-house breeding was conducted using homozygous pairings for Oprk1*^lox/lox^* mice and KtdT mice, and heterozygous pairings for KOR-iCre mice to produce homozygous mice.

Adult male and female mice used in an age (2-8 months old)-matched manner in this study. Animals were group-housed under standard laboratory conditions and kept on a 12 h day/ 12h night cycle (lights on at 6:00 a.m.). Mice were maintained in accordance with the National Institutes of Health Guide for the Care and Use of Laboratory Animals. All methods used were preapproved by the Institutional Animal Care and Use Committee at Temple University.

### Perfusion of mice with 4% paraformaldehyde (PFA) and cryoprotection

Mice were deeply anesthetized with sodium pentobarbital (200-300 mg/kg, i.p.) and perfused transcardially with 20 ml 0.1 m phosphate buffered saline (1× PBS; pH 7.4) followed by 20 ml 4% PFA solution in 0.1 m phosphate buffer (1× PB; pH 7.4). Brains were dissected, postfixed in 4% PFA in 1× PB overnight, and then placed in 30% sucrose for up to 72 h for cryoprotection.

### *In situ* hybridization (ISH) of KOR, vGluT2, and vGluT1

ISH was conducted as described previously [38]. C57BL/6N mice were perfused with 4% PFA and brains were post-fixed and cryoprotected as described above. Brains were frozen in O.C.T. and sectioned at −18°C at 14 μm with a cryostat (Leica CM3050S), mounted onto Superfrost Plus slides and kept at −80°C for less than three months. Advanced Cell Diagnostics (ACD) RNAscope Technology was used for ISH with three mouse probes for KOR (Mm-Oprk1-C2, 316111-C2), vGluT1 (Mm-Slc17a7, 416631) and vGluT2 (Mm-Slc17a6-C3, 319171-C3). Tissue sections were thawed at room temperature briefly, washed with 1× PBS and subsequently processed per the ACD’s protocols. Tissue sections were then permeabilized with solutions from the pretreatment kit and incubated with protease for 30 min and hybridization probes for another 2 h at 40°C.

### Immunohistochemistry (IHC) of KOR-tdT in mouse brain sections

KOR-tdT mice were perfused with 4% PFA and brains were post-fixed and cryoprotected as described above. Brains were frozen in O.C.T. and sectioned with a cryostat (Leica CM3050S) at a thickness of 30 μm at −18°C and placed in 10 mM PBS (8.2 mm Na_2_HPO_4_, 1.8 mm KH_2_PO_4_, NaCl 134 mm, and KCl 2.7 mm; pH 7.5) plus 0.05 NaN3% for short-term storage at 4°C. Sections were rinsed with 10 mM PBS 5 × 5 min, blocked for one hour at room temperature with the blocking buffer (5% normal goat serum, 0.1 m glycine, and 0.3% Triton X-100 in 10 mm PBS). Sections were incubated with rabbit anti-RFP (1:2000, Rockland, 600-401-379) in the staining buffer (3% BSA and 0.3% Triton X-100 in 10 mm PBS) at 4°C overnight and washed 5 × 5 min with 10 mm PBS. Sections were then incubated with Alexa Fluor 594-conjugated goat anti-rabbit IgG (Invitrogen, A11012) (1:1000) overnight at 4°C and washed 5 × 5 min with 10 mm PBS. Sections were subsequently mounted on fluorescence-free glass slides with VECTASHIELD containing DAPI and placed at 4°C for storage for up to two months. Sections were examined under a fluorescence microscope (Keyence, BZ-X800).

### Stereotaxic viral injections into aPVT of the mouse brain

Mice were anesthetized with ketamine (72 mg/kg) and xylazine (7.2 mg/kg) i.p. and placed in a stereotaxic apparatus (Kopf). Viral vectors were injected into the aPVT of mouse brains using the coordinates (relative to the bregma) of AP −0.50, ML 0.0, DV - 3.45 as modified from those of Paxinos and Franklin [33]. Two methods were used: iontophoresis and micropump injections. Iontophoresis was performed using a Stoelting Midguard current source device. The parameters were 5 µA, alternating 7s, 5min with 2 min delay before and after current on/off. Micropump injections (50–75 nl) were administered via a 35G needle attached to a UMP3 micropump (World Precision Instruments).

### Anterograde tracing

We used KOR-iCre mice and two different Cre-dependent anterograde tracers: AAV8-hSyn-DIO-mCherry and AAV1-FLExloxP-mGFP-2A-synaptophysin-mRuby.

AAV1-FLEx-mGFP-2A-synaptophysin-mRuby: mGFP is membrane-associated GFP due to its palmitoylation and synaptophysin-mRuby is used as a marker of presynaptic sites. On day 0, AAV1-FLExloxP-mGFP-2A-synaptophysin-mRuby was stereotaxically injected by Iontophoresis or using a micropump into the aPVT of KOR-iCre mice. Starting from day 14, mice were administered tamoxifen (3 mg/mouse) by gavage daily for 5 days to induce Cre expression. One month later, mice were perfused with 4% PFA and brains were, post-fixed, and cryoprotected as described above. Brains were sectioned coronally at −18°C at 30 μm and sections were mounted sequentially on 24 gelatin-subbed slides. IHC was performed with goat anti-GFP (1:2000, Rockland 600-101-215) followed by Alexa Fluor 488-donkey anti-goat IgG (1:1000, invitrogen A11055) and rabbit anti-RFP antibodies (1:2000, Rockland 600-401-379) followed by Alexa Fluor 594-donkey anti-rabbit IgG (1:1000, invitrogen A21207). Sections were imaged with a Keyence BZ-X800 fluorescence microscope (4X, 10X, 60X objective).

AAV8-hSyn-DIO-mCherry: Stereotaxic viral injections, tamoxifen treatment, perfusion and sectioning were performed as described above. IHC was performed with rabbit anti-RFP antibodies (1:2000, Rockland 600-401-379) followed by Alexa Fluor 594-donkey anti-rabbit IgG (1:1000, invitrogen, A11012).

### KOR conditional knockdown (KOR cKD): injections of viral vectors and cKD confirmation

AAV-eGFP-Cre or AAV-eGFP virion was injected into aPVT of oprk1^lox/lox^ mice as described above. At least one month after viral injections, the mice were tested for a battery of behavioral tests (see below). Subsequently, mice were euthanized by decapitation following completion of behavioral testing. Brains were rapidly removed and immediately froze in isopentane on dry ice. Frozen brains were cut at 20 μm to obtain coronal sections at −18°C, which were thaw-mounted onto gelatin-subbed slides. The sections were first scanned for eGFP expression on a Keyence slide scanner to verify viral injection site and to determine viral spread. Sections were then stored at −80 °C for autoradiography. The sections were processed for autoradiography of [^3^H]U69,593 binding to KOR which allowed evaluation of KOR cKO efficiency and accuracy by aligning up ‘multiplexed’ fluorescent viral expression images with KOR binding signals from the same sections.

### KOR autoradiography

The experiments were performed per our published procedures [43, 44]. Brain sections were taken out of −80°C, thawed at RT for 1h, washed once with 50 mM Tris-HCl buffer (pH 7.4) followed by ∼5 nM [^3^H]U69,593 in 50 mM Tris-HCl buffer with or without 10 μM naloxone at room temperature for 1 h. Slides were then rinsed three times with 50 mM Tris-HCl buffer at 4°C and once with deionized water and then quickly dried with cold air and dried further in desiccator overnight. Sections were then exposed to ^3^H-sensitive phosphor screens (Cytiva) for about four weeks and images on the screens were captured with a Cyclone Storage Phosphor Scanner (Packard Bioscience). Images were inputted into NeuroInfo for alignment of brain regions with the mouse brain atlas of Allen Brain and density was quantified accordingly.

### Mouse behavioral tests

A battery of up to six of the following seven behavioral tests was conducted in the same cohort of Oprk1^lox/lox^ mice beginning at least 1 month after AAV-eGFP-Cre or AAV-eGFP injection. No cohort was used for both morphine withdrawal and fear conditioning experiments. Depending on the cohort, mice underwent up to six tests in the sequence described below, with a 3–7-day washout period between tests to minimize carryover effects.

**Conditioned place aversion (CPA)** was assessed using a modified version of our previously described conditioned place preference procedure [45], utilizing two-chamber boxes as detailed in our previous work [46–48]. The experimental timeline spanned five days. On Day 1, mice underwent a 15-minute pre-test to record baseline chamber preference. On days 2-4, mice were pretreated with (morning / afternoon) saline / saline (10 ml/kg, s.c.) or saline / U50,488H (2 mg/kg, s.c.) 10 min before each conditioning session, in which mouse was confined to one chamber for 30 min. A biased experimental design was implemented due to constraints in available mouse numbers following breeding and surgical procedures. On Day 5, a final 15-minute post-test measured the time spent in each chamber.

**Elevated plus maze (EPM) test** was performed as we previously described [49, 50], with minor modifications, using a mouse EPM apparatus (San Diego Instruments, Inc.). Testing was conducted under red lighting conditions. At the start of each session, mice were placed in the center of the maze facing a closed arm. Exploratory behavior was videotaped, and entries into the open arms, closed arms, and center were recorded during the first 5 min of EPM exposure. An entry into an arm or the center was defined as placement of all four paws into that compartment; otherwise, the animal was considered to remain in the previous compartment. The maze was cleaned and dried between trials. Anxiety-like behavior was assessed by the percentage of open arm entries or the percentage of time spent in the open arms, calculated relative to total arm entries or total time spent in open and closed arms, respectively. Closed arm entries were used as measures of locomotor activity.

**Compound 48/80 scratching test** was performed as we described [46, 48], which was based on the procedures of Wang, et al. [51]. Briefly, after acclimation, mice were injected with saline (10 ml/kg, s.c.) or U50,488H (5 mg/kg, s.c.) 20 min before administration of compound 48/80 (50 μg, s.c.) into the nape, then the bouts of scratching were counted for 30 min.

**Forced swim test (FST)** was performed following our published protocol [52]. Mice were placed for 6 min in a cylindrical tank (46 cm height × 20 cm diameter) filled to a depth of ∼15 cm with water maintained at 23–25 °C. Swim sessions were videotaped for subsequent analysis. Videos were scored by experimenters blinded to treatment conditions. The duration of immobility—defined as minimal movements necessary to remain afloat—was quantified during the final 4 min of the swim session.

**Fear conditioning** was conducted as previously described by Kutlu and colleagues [53]. <u>Day 1 – Training</u>: Mice were placed in a conditioning chamber, and baseline freezing was assessed for 120 s. Animals then received two conditioned stimulus (CS; white noise, WN)–unconditioned stimulus (US; foot shock, FS) pairings, in which a 30 s CS co-terminated with a 2 s, 0.57 mA foot shock. A 120 s intertrial interval followed the first WN–FS pairing. After the second pairing, mice remained in the chamber for an additional 120 s before being returned to their home cages. <u>Day 2 – Contextual Fear Memory Test</u>: Mice were returned to the original conditioning chamber (Context A) to assess contextual fear memory. Freezing behavior was recorded for 4 min in the absence of any stimuli. <u>Day 3 – Cued Fear Memory Test</u>: To assess cued fear memory, mice were placed in the same chamber modified to a novel context (Context B) by altering the floor and wall textures. Freezing was measured for 3 min in the absence of the CS (pre-CS), followed by 3 min during presentation of the white noise CS.

**Morphine withdrawal** was precipitated by naloxone using a protocol adapted from our previous study [48]. Mice received twice-daily subcutaneous (s.c.) injections of morphine (10:00 and 18:00) for 5 consecutive days with escalating doses (day 1: 20 mg/kg; day 2: 40 mg/kg; day 3: 60 mg/kg; day 4: 80 mg/kg; day 5: 100 mg/kg). On the morning of day 6, mice received an additional injection of morphine (100 mg/kg) or saline. Two hours later, withdrawal was precipitated by a s.c. injection of naloxone (10 mg/kg). Mice were then placed in a transparent cylindrical chamber (50 cm height × 12 cm diameter) and were videotaped for 30 min. The total number of jumps was recorded, with most jumping behavior occurring within the first 5–10 min of observation. Other somatic signs of withdrawal commonly observed in rats were minimal or absent, including wet dog shakes, paw tremors, sniffing, teeth chattering, ptosis, and piloerection.

**Acetic Acid Writhing Test** was carried out according to our published procedures [48]. Mice were individually habituated in the observation chambers for at least 1 h and then pretreated s.c. with either saline or U50 (5 mg/kg). Twenty min later, acetic acid (0.6%) was injected intraperitoneally (10 µl/g, i.p.). The number of writhes (abdominal stretches) was recorded for 20 min continuously.

**Statistical analyses** were performed using GraphPad Prism version 10.3.1 (GraphPad Software Inc., La Jolla, CA). All data are expressed as the mean ± S.E.M. Two-way ANOVA or unpaired t tests were used. Individual group comparisons were performed with Sidak’s multiple comparisons test following two-way ANOVA. P < 0.05 was considered to indicate a statistically significant difference.

## RESULTS

### KOR mRNA is co-expressed with vGluT2 mRNA in the majority of PVT neurons

In brains of male C57BL/6N wildtype mice, RNAscope ISH revealed that KOR mRNA was present in many PVT neurons (Fig. 1A), consistent with the results in the Mouse Brain Atlas of Allen Brain Institute [54]. In addition, ISH also revealed that vGluT2 mRNA, but not vGluT1 mRNA, was expressed in the PVT, in agreement with the finding that the PVT is composed predominantly of vGluT2(+) excitatory projection neurons [15]. When the ISH images of KOR mRNA, vGluT2 mRNA, and vGluT1 mRNA were merged, the results showed that KOR mRNA was expressed in most of these vGluT2 neurons and there was no overlap between KOR mRNA and vGluT1 mRNA (Fig. 1A).

**Figure 1.**
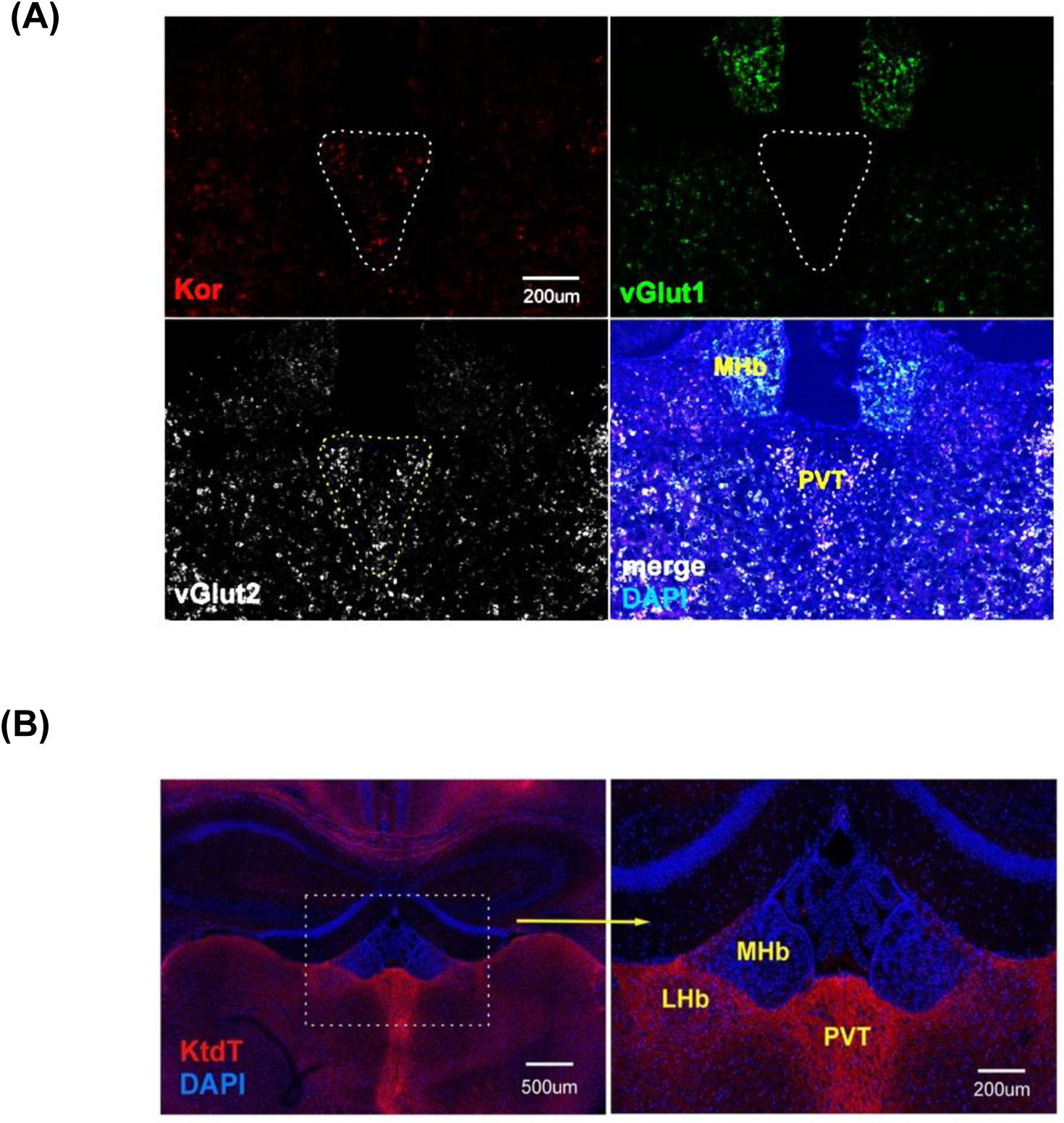
KOR is expressed in PVT. **(A) KOR mRNA is co-localized with vGluT2 mRNA in most PVT neurons.** The PVT boundary is marked with white dash lines. KOR mRNA (red) and vGluT2 (white) are present in the PVT and pink color in the merged image indicates their colocalization in the PVT. vGluT1 mRNA (green) is not present in the PVT. The experiment was performed on two mice with similar results. Abbreviations: MHb: medial habenula; PVT: paraventricular nucleus of thalamus. **(B) KOR protein is abundantly expressed in PVT.** Distribution of KtdT is demonstrated in a coronal section (50 μm thickness) containing the PVT of homozygous KtdT mouse brain. Images were captured with a confocal microscope at −0.94 mm to the bregma. Note that the PVT expresses a high level of KtdT. Experiments were performed on two mouse brains with similar results.

### PVT expresses a high level of KOR protein

We previously generated a knock-in mouse line expressing KOR fused at the C terminus with the fluorescent protein tdTomato (KtdT) [38]. IHC experiments using antibodies against tdTomato demonstrated that KOR protein is abundantly expressed in PVT in the KtdT mice (Fig. 1B).

### Anterograde tracing of KOR(+) neurons in the PVT to determine their efferent projection regions

Anterograde tracing of the aPVT was carried out with AAV8-hSyn-DIO-mCherry injected into the aPVT of KOR-iCre mice (Fig. 2 upper panel). Following IHC against mCherry and imaging, selected images were assembled into a montage (Fig. 2 upper panel) and a video (Supplemental video 1). In addition, images were loaded into the NeuroInfo software and re-constructed into 3-D images, and a video of a 3-D image is shown as Supplemental video 2. The results show dense projections to the NAc, Amyg, RT, and olfactory tubercule (OT) and less dense projections to the BNST, claustrum (CLA), endopiriform nucleus, and prefrontal cortex. It is noteworthy that AAV8 has limited trans-synaptic spread.

**Figure 2.**
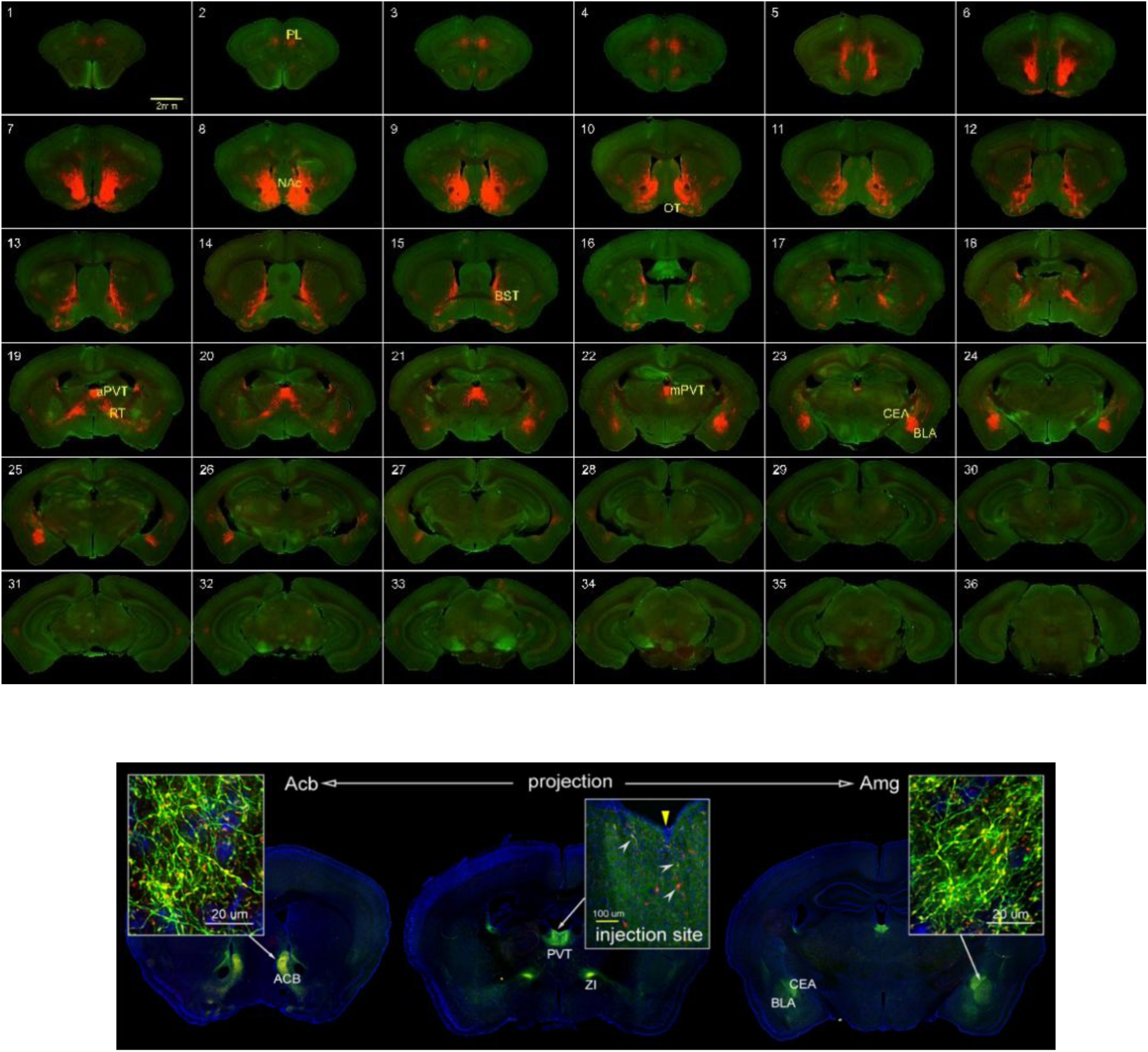
Determination of efferent projection regions of PVT KOR(+) neurons by anterograde tracing in KOR-iCre mice. **Upper Image.** Representative montage at 4X showing projection sites. The NAc, olfactory tubercle, CeA, BLA, BNST, and RT received dense projections from PVT KOR+ neurons. The prelimbic cortex (PL), claustrum (CLA), periaqueductal grey (PAG) (not shown) and VTA (not shown) received moderate projections. **Lower Image.** The coronal section in the middle shows the injection site [the PVT] and two projection sites, the reticular nucleus of the thalamus and stria terminalis (ST). The white arrowheads in the inset point to some (not all) KOR(+) cells (yellow or red). The yellow reverse triangle indicates the path of injection. The coronal sections on the left and right reveal that the medial shell of NAc, CeA and BLA, respectively, received dense projections from KOR(+) cells in the PVT. The insets show GFP+ nerve fibers and GFP+/synaptophysin mRuby+ puncta (vesicles-containing terminals). Blue: DAPI.

The 3-D video (Supplemental video 2) revealed that aPVT projections first targeted the ventral RT, then the NAc, and ultimately project to distal regions including the amygdala. This projection sequence supports earlier findings of robust axonal collateralization in forebrain targets. Furthermore, it corroborates the observation that spatially intermixed PVT neurons simultaneously collateralize to the NAc, BNST, and CeA [16, 17].

Anterograde tracing was also performed with Cre-dependent AAV1-FLExloxP-mGFP-2A-synaptophysin-mRuby injected into aPVT of KOR-iCre mice (Fig. 2, lower panel). Membrane-targeted GFP (mGFP) localizes to the neuronal membrane via palmitoylation, while synaptophysin-mRuby labels putative presynaptic sites (see the insets of the lower panel of Fig. 2. Dense projections from aPVT KOR(+) neurons were observed in the NAc, CeA, BLA, BNST, and RT. Moderate projections were detected in the PFC and CLA.

### Cre-dependent conditional KOR knock-down (KOR cKD) in aPVT revealed by receptor autoradiography

At least one month after AAV-eGFP-Cre or AAV-eGFP injections into the aPVT of oprk1^lox/lox^ mice, animals were euthanized, and brains were collected for KOR autoradiography combined with eGFP fluorescence scanning. KOR autoradiography was performed with ∼5 nM [^3^H]U69,593 with or without 10 μM naloxone. eGFP fluorescence images confirmed accurate targeting of the injection into the aPVT and autoradiography images showed substantial decreases in KOR, indicating efficient Cre-dependent KOR cKD in the aPVT following AAV-eGFP-Cre, but not after control AAV-eGFP injection (Fig. 3).

**Figure 3.**
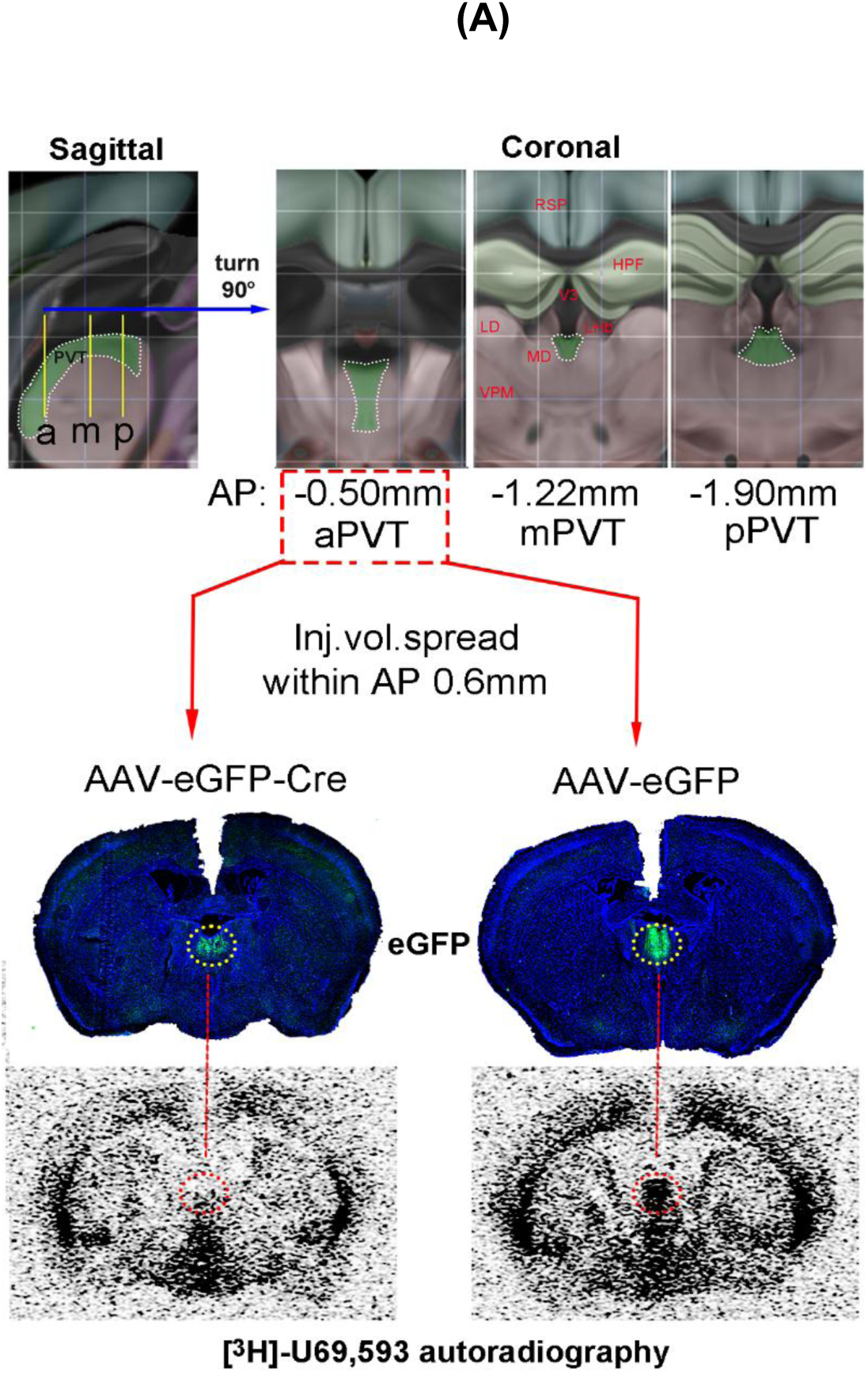
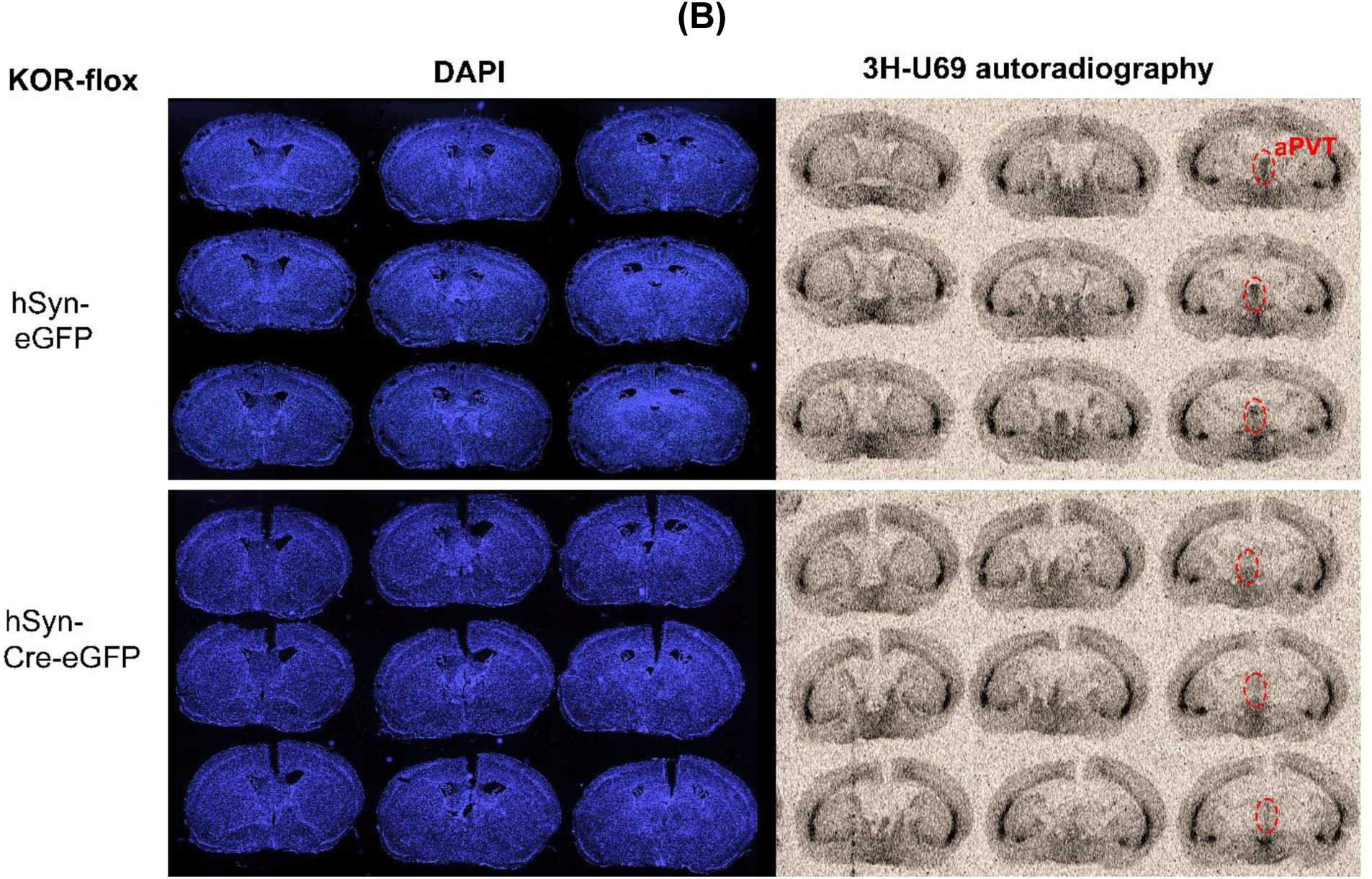
Conditional KOR Knock-down (KOR cKD) in aPVT of oprk1^lox/lox^ mice. **(A)** AAV-eGFP-Cre or AAV-eGFP was injected into oprk1^lox/lox^ mice. Injections targeted the aPVT (AP −0.50, ML 0.0, DV −3.40). At least one-month post-injection, mice underwent CPA and additional behavioral testing, after which fresh brains were collected, sectioned at 20 µm, and mounted for autoradiography. Sections were first scanned for eGFP expression using a Keyence slide scanner to verify injection sites and regional specificity, and then to assess KOR cKD efficiency by aligning viral expression with KOR binding signals. **(B)** more representative data on KOR cKD in aPVT.

### The effects of KOR cKD in aPVT on mouse behaviors

About one month after Oprk1^lox/lox^ mice were injected with AAV-eGFP-Cre or AAV-eGFP, the mice underwent a battery of behavioral tests.

Following the conclusion of behavioral tests, mice were sacrificed and brains were removed. Frozen brain sections were obtained for verification of viral injection sites and for KOR receptor autoradiography. The data obtained from mice with off-target injections or insufficient KOR knockdown were excluded from analysis.

#### Morphine withdrawal

Following injections of escalating doses of morphine (20–100 mg/kg) or saline for 5 and 1/2 days, administration of naloxone (10 mg/kg, s.c.) 2 h after the last injection precipitated withdrawal behavior, which was assessed for 30 min. Total jumps were recorded, with most occurring during the first 5–10 min. Other withdrawal signs observed in rats, including wet-dog shakes, paw tremors, sniffing, teeth chattering, ptosis, and piloerection, were rare or absent. Oprk1^lox/lox^ mice injected with AAV-eGFP-Cre exhibited fewer jumps than AAV-eGFP–injected controls (p<0.05) in both male and female mice, indicating that KOR cKD in the aPVT significantly decreased naloxone-precipitated morphine withdrawal–associated jumping (Fig. 4A).

**Figure 4.**
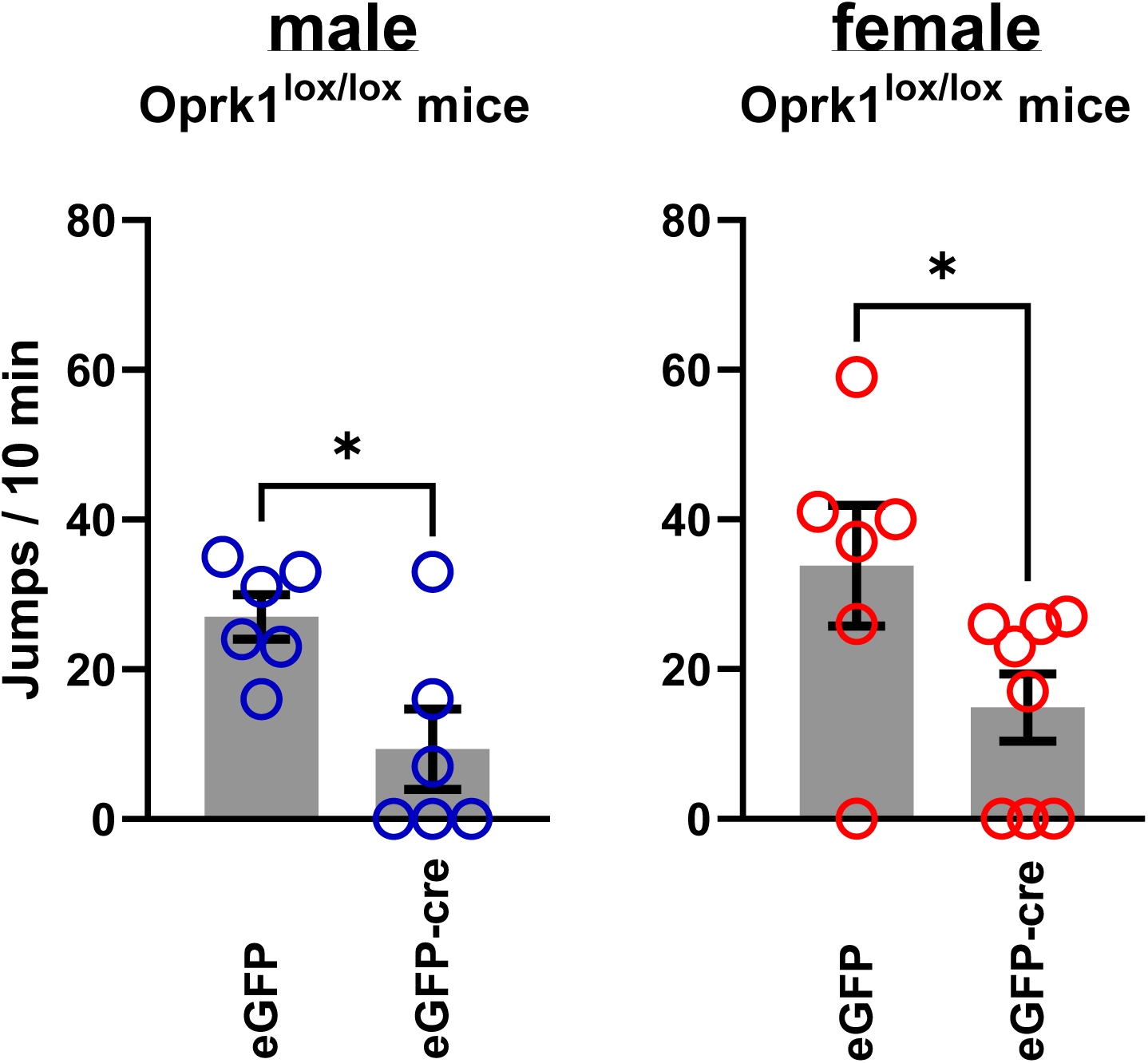
KOR cKD affected mouse behaviors. **A. KOR cKD in aPVT attenuated naloxone-precipitated morphine withdrawal-associated jump in either sex.** Mice underwent twice daily (10:00 and 18:00) treatment for 5 days with escalating doses of morphine via subcutaneous injections (day 1: 20 mg/kg, day 2: 40 mg/kg, day 3: 60 mg/kg, day 4: 80 mg/kg, day 5: 100 mg/kg) or saline. On the morning of day 6, mice received another injection of 100 mg/kg morphine or saline. Two hours later, withdrawal was induced by a subcutaneous injection of 10 mg/kg naloxone. Mice were placed in a 50 cm high, 12 cm diameter transparent cylinder, positioned on white paper, under normal room lighting and observed and videotaped for 30 minutes. Total numbers of jumps were recorded, with the majority of jumping occurring primarily during the initial 5 to 10 minutes. Other withdrawal signs usually observed in rats were minimal, if any, such as wet dog shakes, paw tremor, sniffs, teeth chattering, ptosis, and piloerection. All data are expressed as the mean ± S.E.M (N=6-8). Unpaired t test was used (*p<0.05).

**Figure 4B.**
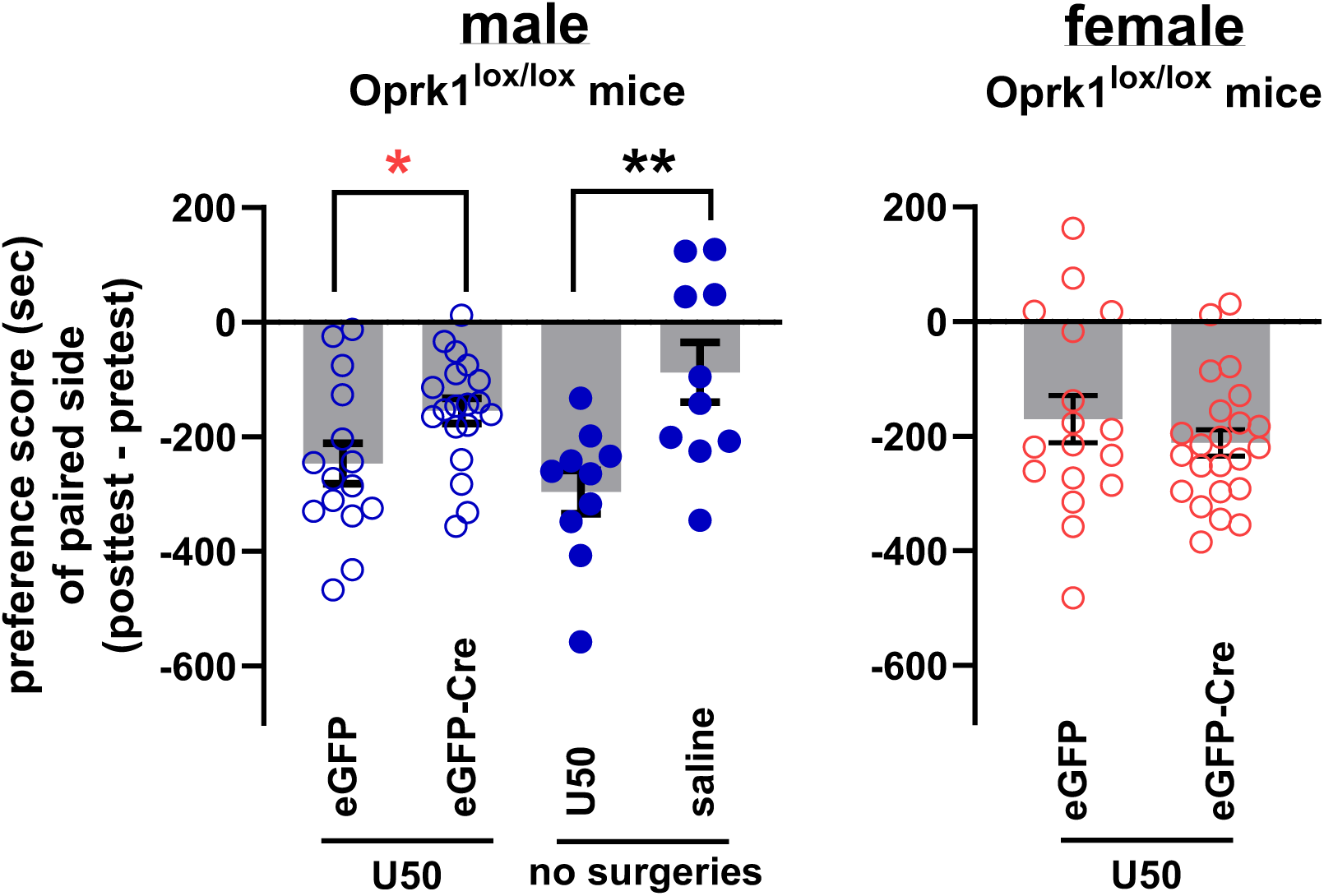
KOR cKD in the aPVT reduced U50-induced conditioned place aversion (CPA) in males but had no effect in females. CPA was performed with two-chamber boxes. On day 1, mice were subject to pre-test for 15 minutes, in which the time each mouse spent in either chamber was recorded. On days 2 to 4, mice were pretreated with (morning/afternoon) saline/saline (10 ml/kg, s.c.) for the “saline” group or saline/U50,488H (2 mg/kg, s.c) for the “U50” group 10 minutes before each conditioning session, in which the mouse was confined to one chamber for 30 minutes. On day 5, the time each mouse spent in either chamber was recorded in the 15-minute post-test. A biased design was used due to the limited number of mice available. All data are expressed as the mean ± S.E.M (N=10-23). Unpaired *t* test was used (*p<0.05, **p<0.01).

**Figure 4C.**
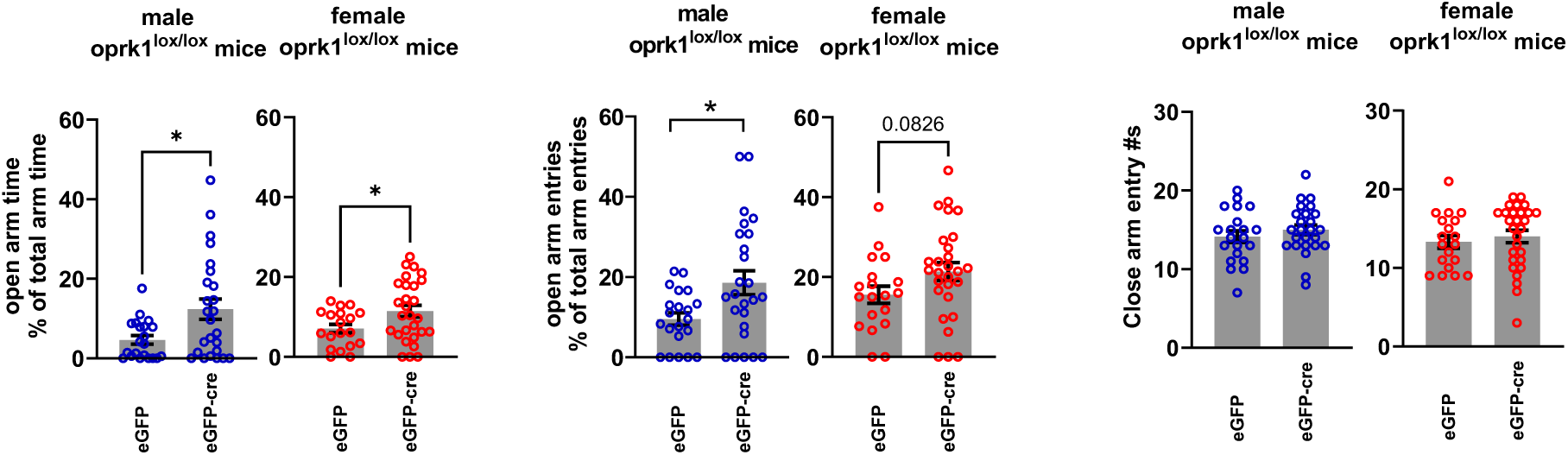
KOR cKD in aPVT of either sex had anxiolytic-like effects in the elevated plus-maze (EPM) test without affecting closed arm entries. EPM test was carried out using a mouse EPM apparatus (San Diego Instruments, Inc.). Mice were tested under red lighting condition. The exploration of mice in the maze was videotaped, and entries into and time spent on open and closed arms were recorded for the first 5 min of EPM exposure. The measures of anxiety are the % of open arm entries and the % of time spent on open arms, both expressed as the % of the total entries into, or time spent on, both the open and closed arms, whereas the numbers of closed arm entries are considered locomotor measures. All data are expressed as the mean ± S.E.M (N=19-28). Unpaired t test was used (*p<0.05).

**Figure 4D.**
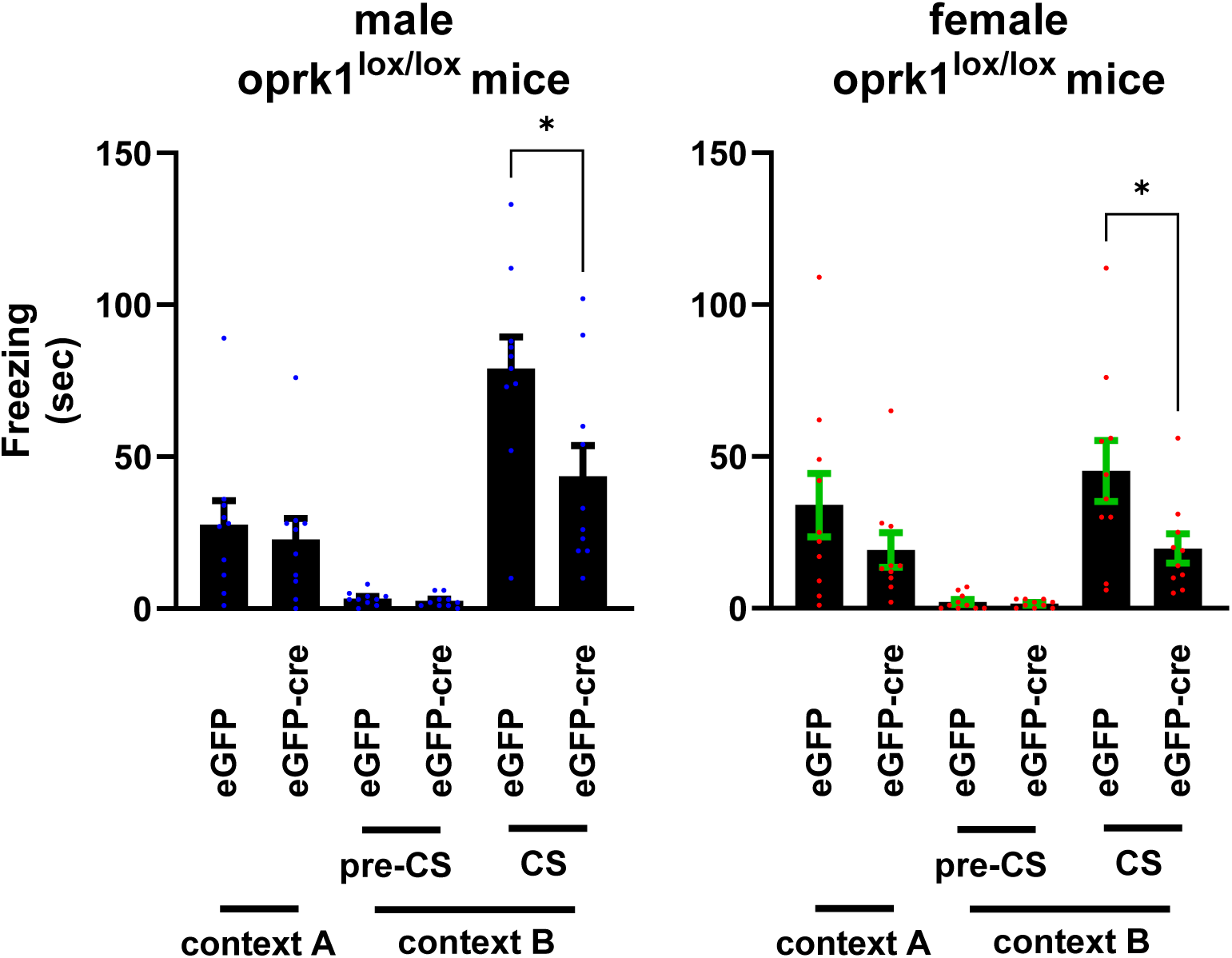
KOR cKD selectively reduced the cue-induced fear after fear conditioning as follows. Day 1 – Training: mice were placed in the conditioning chamber #4 and baseline freezing was assessed for 120 s. Mice then received two CS (white noise (WN))–US [footshock (FS)] pairings in which a 30 s CS co-terminated with a 2 s 0.57 mA FS. After the first WN–FS pairing, there was a 120-s interval. Animals remained in the chamber for 120 s after the second WN–FS pairing and were then removed. Day 2 – Contextual Fear Memory Testing: Mice were first returned to the original chamber (Context A) to assess contextual fear memory, and freezing behavior was recorded for 4 minutes in the absence of any stimuli. Day 3 – Cued Fear Memory Testing: mice were placed in the same chamber modified to Context B (with a plexiglass floor) to assess cued fear memory. In Context B, freezing was measured for 3 minutes without CS (pre-CS), followed by 3 minutes with white noise (CS). All data are expressed as the mean ± S.E.M (N=10). Unpaired t test was used (*p<0.05).

#### U50-induced CPA

CPA was assessed using a two-chamber apparatus. Following a pre-test on day 1, mice underwent conditioning on days 2–4, and CPA was evaluated on day 5. U50 induced significant CPA in non-surgical male Oprk1^lox/lox^ mice compared with saline controls (**p < 0.01) (Fig. 4B). In male Oprk1^lox/lox^ mice, U50-induced CPA was significantly attenuated by AAV-eGFP-Cre injection relative to AAV-eGFP controls (*p < 0.05) (Fig. 4B). In contrast, U50-induced CPA was comparable between AAV-eGFP-Cre– and AAV-eGFP–injected female mice (Fig. 4B). These findings indicate that KOR cKD in the aPVT reduced U50-induced CPA in males but not females.

#### EPM

Mouse exploration on EPM was videotaped for 5 min, and entries into and time spent on open and closed arms were recorded and quantified. In male Oprk1^lox/lox^ mice, AAV-eGFP-Cre injections significantly increased the percentage of open-arm entries and time compared with AAV-eGFP controls (\**p* < 0.05). In females, open-arm time was significantly increased (\**p* < 0.05), and open-arm entries showed a trend toward increase (*p* = 0.0826). Closed-arm entries were unchanged in both sexes, indicating no effect on locomotor activity (Fig. 4C). Overall, KOR cKD produced anxiolytic-like effects in both males and females, as reflected by increased open-arm exploration without changes in locomotion (Fig. 4C).

#### Fear conditioning

Fear conditioning was carried out on Day 1. Contextual testing was performed on Day 2, in which freezing was assessed for 4 min in the original context (Context A) without stimuli. Cued testing was conducted on Day 3, when mice were tested in Context B for 3 min pre-CS and 3 min during CS (white noise). In both male and female Oprk1^lox/lox^ mice, AAV-eGFP-Cre injections significantly reduced CS-induced freezing compared with AAV-eGFP controls, while pre-CS freezing was minimal (Fig. 4D). Contextual freezing did not differ between the two groups (Fig. 4D). Thus, KOR cKD reduced cue-induced fear without affecting contextual fear in either sex.

In addition, male and female Oprk1^lox/lox^ mice injected with AAV-eGFP-Cre or AAV-eGFP were assessed in the following mouse behavioral test. Results demonstrated that KOR cKD in the aPVT did not affect immobility time in the forced swim test (FST) (Supplemental Fig. 2), the anti-scratching effects of U50 (Supplemental Fig. 3), or the antinociceptive effects of U50 in the acetic acid–induced writhing test in both males and females (Supplemental Fig. 4).

## DISCUSSION

Our results indicate, for the first time, that KOR signaling in the aPVT contributes to morphine withdrawal-related, anxiety-like, fear-related, and KOR agonist-evoked aversion-like behaviors. Historically, behavioral studies of the PVT have focused primarily on the pPVT [14], whereas investigations of the aPVT have only recently begun to emerge (e.g., [20–32]). To the best of our knowledge, our results uncover previously unrecognized contributions of the aPVT to the regulation of morphine withdrawal-like behaviors, acting in a manner that appears to differ from the pPVT.

### Efferent projections of KOR(+) neurons in the aPVT

We found that KOR(+) neurons in the aPVT project to (from dense to moderate) the NAc, BNST, RT, BLA, OT, PFC, CeA and CLA,.The aPVT was shown to send efferent projections to (from dense to moderate) PFC, NAc, BLA, LH, BNST, and ventral subiculum [reviewed in [15]]. The relative densities of aPVT KOR(+) neuronal projections are different from those of aPVT projections, likely due to variations in KOR expression levels in neurons projecting to different targets.

### Contribution of the aPVT to morphine withdrawal-like behavior

Our observations on the role of aPVT KOR in morphine withdrawal are reminiscent of the following two studies. Kelsey et al. [55] showed that the selective KOR antagonist nor-binaltorphimine (nor-BNI) decreased both somatic signs and CPA associated with naloxone-precipitated morphine withdrawal in rats [55]. Constitutive KOR gene disruption markedly reduced morphine withdrawal signs in mice [2]. Our findings demonstrate that targeted disruption of postsynaptic KOR in the aPVT specifically is sufficient to achieve a similar effect on the precipitated withdrawal-associated jumping behaviors in mice. We attempted to assess the effect of aPVT KOR cKD on withdrawal-induced CPA in mice; however, this analysis could not be performed because withdrawal-induced CPA was not reliably observed in wildtype mice with our experimental paradigm.

Antagonist-precipitated morphine withdrawal increased c-Fos expression in both aPVT and pPVT neurons in guinea pigs [56] and in pPVT in mice[57]. While previous studies have emphasized the role of pPVT and its downstream circuits in mediating the aversive and physical components of opiate withdrawal [57–59], our findings indicate that KOR activation in aPVT neurons is related to increased withdrawal-associated jumping. Therefore, our study identified a previously unrecognized contribution of the aPVT to morphine withdrawal–associated behaviors and provided causal evidence for a functional role of aPVT KOR signaling in withdrawal behaviors. Since KOR activation suppresses glutamatergic neurons in the aPVT, KOR cKD would likely disinhibit aPVT neuronal activity, which in turn exerts anti-withdrawal effects. In contrast, inhibiting pPVT attenuated the withdrawal [57]. Therefore, aPVT and pPVT play divergent roles in opioid withdrawal.

### Contribution of the aPVT to anxiety-like behavior

Given that KOR activation inhibits glutamatergic neurons in the aPVT, removing this inhibition by KOR cKD likely disinhibits aPVT neuronal activity, which in turn appears to produce anxiolytic effects. Our finding, to our knowledge, constitutes the first evidence implicating the aPVT in anxiety-like behaviors in normal animals. Optogenetic activation of the aPVT–NAc pathway did not alter anxiety-like behaviors in the EPM or light–dark box tests in mice [61], likely because other aPVT projections are involved.

Microinjection of the GABA_B_ and GABA_A_ receptor agonists baclofen and muscimol into the aPVT had no effect on open-arm time and open-arm entries in the EPM test [62]. However, the aPVT has recently been implicated in anxiety regulation in animals with chronic visceral pain [20]. Briefly, adult offspring of prenatal maternal stress-exposed mice exhibited visceral hypersensitivity and anxiety-like behaviors. In these mice aPVT-BLA circuit regulated anxiety. Increased expression of *Cacna1e* (encoding the alpha-1E subunit of Cav2.3 voltage-gated calcium channels) in the aPVT potentiated both visceral pain and anxiety [20].

The involvement of aPVT KOR in anxiety-like behaviors observed here aligns with previous reports (see [8] for review) that KOR antagonists or KOR gene deletion produces anxiolytic effects. Long-acting KOR antagonists reduce anxiety-like behavior in the EPM, fear-potentiated startle, novelty-induced hypophagia (NIH), and defensive burying assays [63–66]. The short-acting KOR antagonists zyklophin and LY2444296 showed anxiolytic-like effects in the NIH test, but not in the EPM test [50]. In addition, anxiolytic-like phenotypes have been observed after ablation of KOR from brain dopamine neurons or deletion of prodynorphin [44, 67]. Notably, it is likely that the KOR(+) neurons in the aPVT projecting to amygdala is involved in anxiety-like behavior. These findings suggest that KOR signaling may contribute to the acquisition and/or expression of anxiety-like behaviors. However, mice with constitutive global KOR knockout showed no differences from the wildtype in measures of anxiety-like behavior [2].

### KOR in aPVT contributed to cued, but not contextual fear

We found that cKD of KOR in the aPVT attenuated cued fear-associated freezing. As KOR activation inhibits aPVT neuronal activity, one possible mechanism is that endogenous dynorphin release during cued fear recall suppresses aPVT neuronal activity via KOR to facilitate fear expression. KOR cKD eliminates this inhibition, resulting in enhanced aPVT activity and consequent suppression of cue-induced fear expression. Notably, KOR cKD in aPVT attenuated cue-, but not context-, induced fear expression. Chen et al.[68] found that glutamatergic neurons in the aPVT were recruited during memory recall of cued fear, as indicated by increased c-Fos expression. Chemogenetic inhibition of aPVT neurons increased, whereas optogenetic activation of these neurons decreased freezing in both contextual and cued fear recall tests. Such modulations by aPVT neurons are opposite to those by pPVT neurons [68]. Therefore, KOR cKD in the aPVT is likely to remove a specific inhibitory modulatory mechanism that may be selectively engaged during discrete cue recall. This raises the possibility that dynorphin–KOR signaling gates amygdala-related cue processing within the aPVT, while contextual fear expression relies on broader hippocampal–thalamic circuitry that is less modulated by KOR tone. The latter is consistent with our finding that aPVT KOR(+) neurons project to the amygdala, but not to the hippocampus.

The involvement of aPVT KOR in mediating the cued fear-like behavior observed here is consistent with previous reports implicating KOR signaling in fear responses. Systemic administration of long-acting KOR antagonists such as norBNI or JDTic or infusions of JDTic into the BLA or CeA reduced cued fear in the fear-potentiated startle paradigm [64, 65]. It is noteworthy that this is the first report that aPVT KOR is at least in part responsible for cue-induced fear. It is likely that the projection from aPVT to amygdala accounts for this behavior.

### The role of aPVT KOR in KOR agonist-induced aversion

Our findings demonstrate that selective disruption of postsynaptic KOR within the aPVT attenuates U50,488H-induced CPA in a sex-specific manner, with effects observed in male mice only. These results suggest that aPVT KOR signaling contributes to the encoding or expression of aversive motivational states but is not the sole substrate for KOR-mediated aversion. The mechanisms underlying the male-specific effect are likely complex and remain to be elucidated.

KOR agonists have been demonstrated to induce CPA [69, 70]. Shippenberg et al. showed that this was mediated by reduced NAc dopamine via inhibition of DA release from VTA terminals [71–73], whereas Chavkin et al. found CPA persisted in dopamine-deficient mice and depended on a DRN→NAc serotonin pathway [74, 75]. aPVT sends dense projections to the NAc to innervate MSNs [61], which are also innervated by the VTA DA neurons and DRN 5-HT neurons. It is likely that modulation of the MSNs by the aPVT projections contributes to aPVT KOR-induced CPA. This is the first recognition that aPVT KOR is involved in producing KOR agonist-induced CPA. Collectively, these findings suggest that the aPVT can modulate mesolimbic dopamine function indirectly via downstream targets such as the NAc.

### Paraventricular thalamus circuits in visceral pain processing

Global KOR knockout has been shown to enhance sensitivity to visceral pain induced by acetic acid in the writhing test [2]. KOR cKD in the aPVT did not affect U50,488H-induced antinociception in the acetic acid writhing test, excluding KOR in the aPVT as an important player in relieving visceral pain. In contrast, pPVT has been shown to play a significant role in relieving visceral painVisceral nociceptive stimulation activates the pPVT and suppressing activity in the pPVT significantly reduced visceral pain–related behaviors [76].

### Concerns about conducting pathway-specific studies

Whether KOR signaling within specific aPVT projection pathways, such as the aPVT–amygdala or aPVT–NAc circuits, mediate opioid withdrawal-associated behavior, anxiety, cued fear memory, and aversion remains an interesting and important question. However, pathway-specific approaches are complicated by axonal collateralization among aPVT projections (Supplemental video 2), which occurs at moderate to high levels across extended forebrain targets [16, 17], and will complicate interpretations of the results. It has been observed that PVT neurons send collateral projections to the NAc, BNST and CeA and that these neurons are intermixed in the aPVT and pPVT [16, 17]. The extent of collateralization varies depending on neuronal subpopulations and the specific regions of the amygdala and NAc they innervate [16, 17].

### Opioid receptors in the aPVT and their roles in behaviors

This study provides the first evidence that KOR in the aPVT regulates behaviors. It is also only the second study to examine the involvement of an opioid receptor type in the aPVT in behaviors. Previously, mu opioid receptor (MOR) signaling in the aPVT has been shown to regulate defensive behaviors elicited by a conditioned fear stimulus via pharmacological manipulation [77].

### Postsynaptic KOR in the aPVT regulated behaviors

The behavioral roles of receptors in the PVT have typically been investigated using pharmacological approaches rather than direct manipulation of receptors in PVT neurons. Such approaches are less selective because agonists and antagonists act on both pre- and post-synaptic receptors. In this study, we used a KOR cKD strategy to selectively remove KORs from aPVT neurons while preserving presynaptic KORs, enabling precise investigation of the postsynaptic contribution of aPVT KOR signaling to behavior.

## Conclusion

Our results reveal for the first time that KOR in the aPVT neurons and their projections critically regulates morphine withdrawal-, anxiety-, cued fear-, and KOR agonist–evoked aversion-like behaviors. These findings uncover previously unrecognized roles for the aPVT. Overall, these data identify the aPVT as a key thalamic hub for KOR-mediated regulation of motivational and affective states and suggest it as a potential target for interventions in opioid withdrawal-, anxiety-, and fear-related disorders.

## Acknowledgement

NIH grants R01DA056581 and P30DA013429 (LYLC). We thank Dr. Munir Gunes Kutlu, PhD (Lewis Katz School of Medicine, Temple University) and his lab members Oyku Dinckol and Noah Wenger, for equipment and technical support related to the fear conditioning experiments.

## Abbreviations

ACD: Advanced Cell Diagnostics
Amyg: amygdala
ANOVA: analysis of variance
aPVT: anterior paraventricular nucleus of the thalamus
BLA: basolateral amygdala
BNST: bed nucleus stria terminalis
BSTDL: dorsolateral BNST
CeA: central amygdala
CeL: central amygdala lateral division
cKD: conditional knockdown
CLA: claustrum
CPA: conditioned place aversion
CS: conditioned stimulus
EPM: elevated plus maze
FS: foot shock
FST: forced swim test
GPCRs: G protein-coupled receptors
IHC: immunohistochemistry
i.p.: intraperitoneally
ISH: in situ hybridization
KOR: kappa opioid receptor
KOR-iCre: a mutant mouse line custom generated by Cyagen Co. that expresses KOR linked to CreERT2 via a 2A peptide
KOR cKD: KOR conditional knockdown
KtdT: a knock-in mouse line expressing KOR fused at the C terminus with the fluorescent protein tdTomato
mGFP: membrane-targeted GFP
MHb: medial habenula
MOR: mu opioid receptor
NAc: nucleus accumbens
NAcSh: nucleus accumbens shell
NIH: novelty-induced hypophagia
nor-BNI: nor-binaltorphimine
PAG: periaqueductal gray
PFC: prefrontal cortex
PL: prelimbic cortex
pPVT: posterior paraventricular nucleus of the thalamus
pre-CS: in the absence of the CS
PVT: paraventricular nucleus of the thalamus
RT: reticular nucleus of the thalamus
s.c.: subcutaneously
snRNA-seq: single nucleus RNA sequencing
ST: stria terminalis
U50: U50,488H
U50,488H: U50,488 methanesulfonate
US: unconditioned stimulus
VTA: ventral tegmental area
WN: white noise
WT: wildtype
ZI: zona incerta

## Supplemental Figures and Legends

**Supplemental Figure 1.**
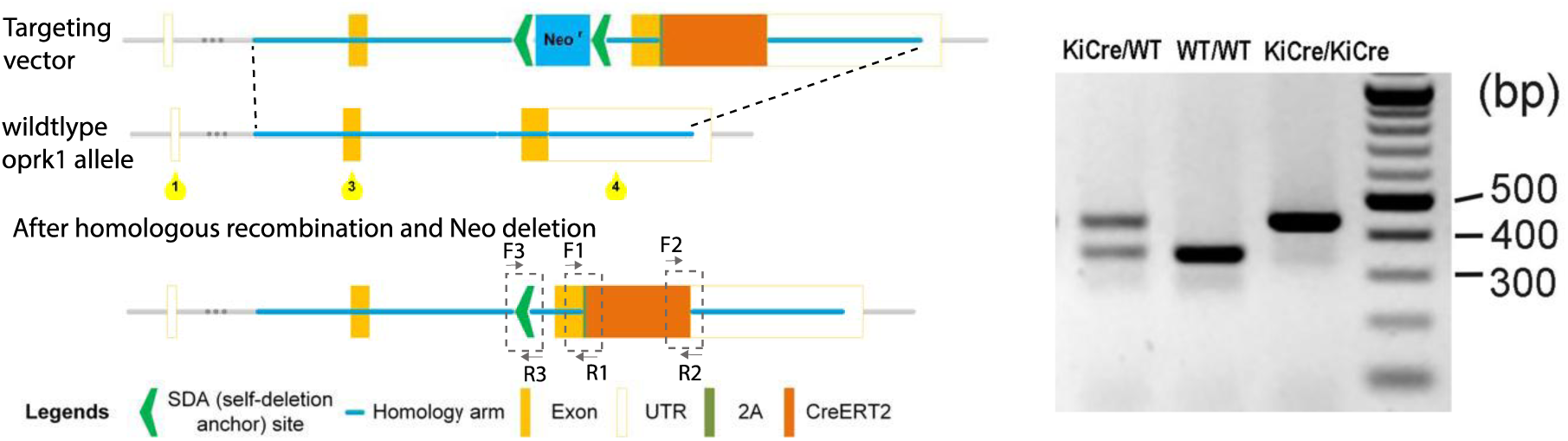
Targeting strategy for generation of KOR-iCre mice. *Oprk1* exons 3 and 4, and the floxed neomycin cassette are shown as orange and blue boxes, respectively. Peptide 2A and CreERT2 are indicated by olive green and dark orange boxes, respectively. Knockin was accomplished by homologous recombination and the Neo cassette was self-deleted in germ cells. Genotyping was performed with F3 and R3 primers for detection of the remaining loxP site in the KOR-iCre mutant mice. The PCR products are 436bp and 347bp for the KOR-iCre and wildtype mice, respectively.

**Supplemental Figure 2.**
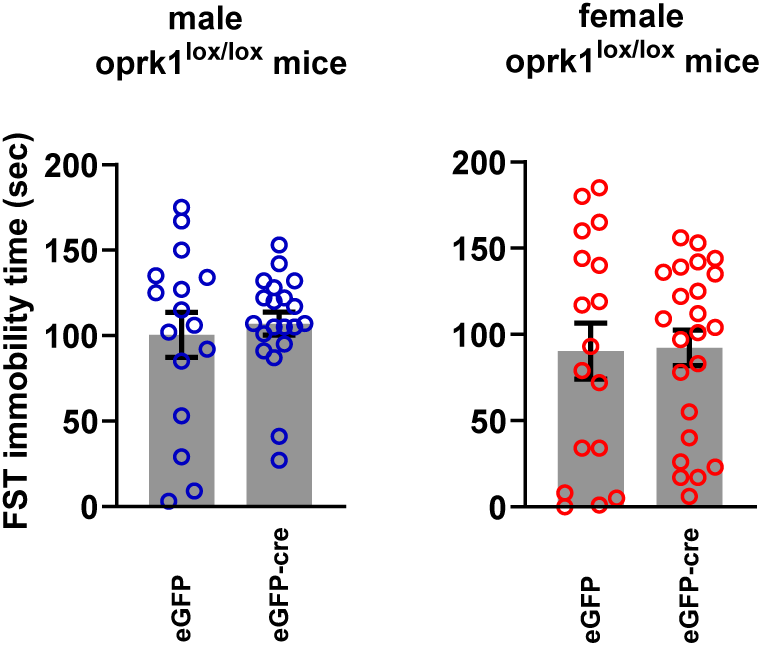
KOR cKD in aPVT did not affect the immobility time in the forced swim test (FST). Mice were placed for 6 min in a cylindrical tank (46 cm tall x 20 cm diameter) of 23–25 °C water filled to a depth of 15 cm. The swim sessions were videotaped and scored later. The FST videos were scored by other researchers in the lab who were blind to the treatments. Duration of immobility (minimum movements necessary to stay afloat) in the last 4-min of the swim was measured. All data are expressed as the mean ± S.E.M (N=17-23). Unpaired t test was used (no significance was found between groups).

**Supplemental Figure 3.**
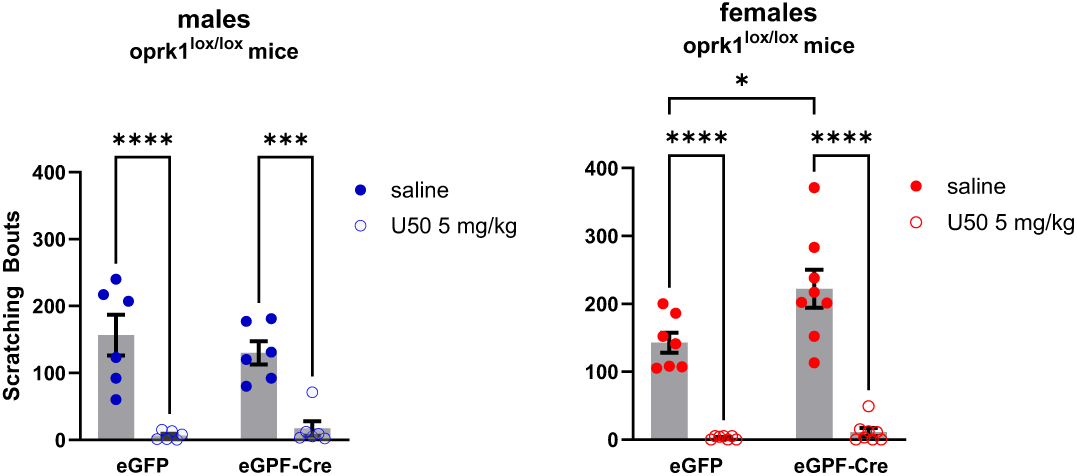
KOR cKD in aPVT did not alter the anti-scratching effect of U50. Mice were allowed to acclimate individual rectangular observation Plexiglass boxes for 1 h and then injected s.c. with either saline or U50,488H (5 mg/kg) in volumes of 0.1 ml/10 g body weight. Twenty min later, animals were injected s.c. with 0.1 ml of compound 48/80 (0.5 mg/ml in saline; 50 μg) into the nape. Compound 48/80 causes release of histamine from mast cells and produces scratching behavior in mice. After 1 min, mice were observed for 30 min and the number of bouts of hindleg scratching movements directed to the neck were counted. All data are expressed as the mean ± S.E.M (N=6-8). Two-way ANOVA was performed followed by the Sidak’s multiple comparisons test (*p<0.05, ***p<0.001, ****p<0.0001).

**Supplemental Figure 4.**
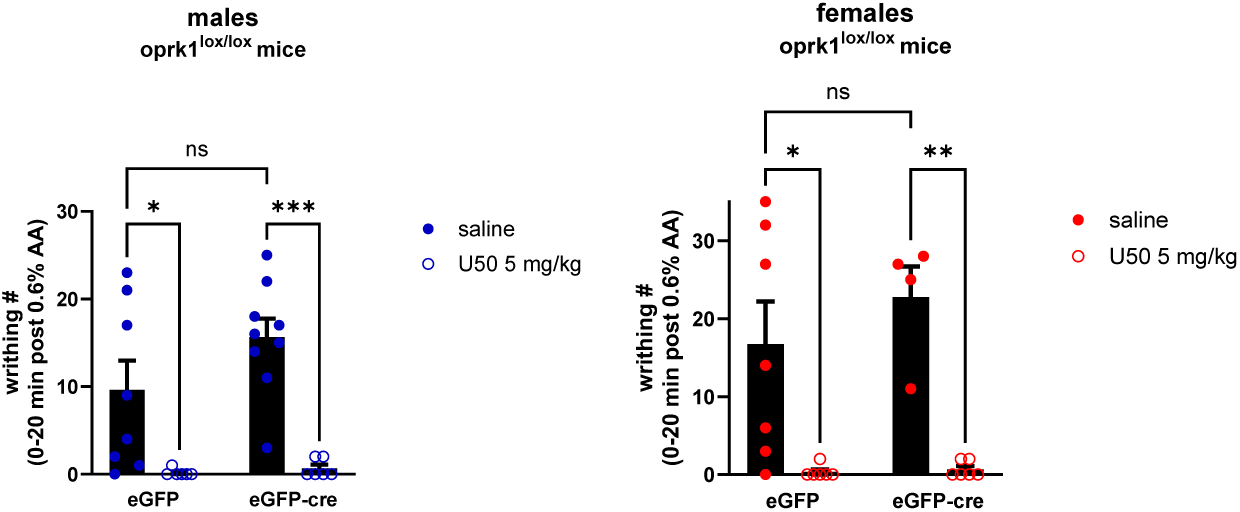
KOR cKD in aPVT did not alter the anti-nociceptive effect of U50 in acetic acid-induced writhing test. Mice were allowed to acclimate to individual rectangular observation Plexiglass boxes for 1 h and then injected s.c. with either saline or U50,488H (5 mg/kg) in volumes of 0.1 ml/10 g body weight. Twenty min later, animals were injected i.p. with 0.6 % acetic acid in saline (0.1 ml/10 g body weight). Mice were observed for 20 min and the number of abdominal stretches were counted. All data are expressed as the mean ± S.E.M (N = 4-9). Two-way ANOVA was performed followed by the Sidak’s multiple comparisons test (*p<0.05, **p<0.01, ***p<0.001).

### Two Supplemental Videos in addition to this PDF file

Supplemental video 1 (selected stacks from Fig. 2 upper panel)

Supplemental video 2 (3-D)

